# Characterizing load-dependent changes in whole-brain activity patterns during an extended N-back task

**DOI:** 10.64898/2026.06.19.733380

**Authors:** Shinya Chiyohara, Tomohisa Asai, Kentaro Hiromitsu, Hiroshi Imamizu

## Abstract

Working memory (WM) is a core cognitive function that supports goal-directed behavior by temporarily maintaining and manipulating information. One of the most widely used paradigms for investigating WM function is the N-back task, and numerous neuroimaging studies have examined load-dependent neural responses using a variety of analytical approaches. However, most previous studies have focused on low-to-moderate load ranges (primarily 0–3-back), and it remains unclear how whole-brain activity patterns reconfigure across a broader range of WM demands, including conditions approaching capacity limits. In the present study, we investigated behavioral performance and whole-brain activity patterns across an extended N-back task ranging from 0-back to 7-back. Behavioral analyses revealed that discrimination sensitivity (d′) decreased nonlinearly with increasing WM load, whereas reaction time (RT) exhibited an inverted-U pattern, peaking at intermediate load conditions. To characterize load-dependent whole-brain activity patterns, we computed relative activation maps by subtracting the participant-wise mean activation map across all conditions from each condition-specific activation map. Spatial similarity analyses with the Yeo 7-network templates revealed that low-load conditions showed relatively high similarity to default mode network (DMN)-related patterns. Similarity to the dorsal attention network (DAN) and frontoparietal network (FPN) was maximal at intermediate load levels, indicating load-dependent changes in network similarity profiles. High-load conditions were characterized by partial re-emergence of DMN-related patterns, accompanied by reduced DAN/FPN similarity. In addition, semantic similarity analysis using Neurosynth-derived semantic maps revealed relatively high similarity to default mode-related and self-referential representations under low-load conditions. Intermediate-load conditions showed strong correspondence with working memory- and executive control-related representations, whereas high-load conditions exhibited increased similarity to salience-, aversive/interoceptive-, and inhibitory-control-related representations. Together, these findings suggest that increasing WM load is associated not merely with stronger activation, but with changes in whole-brain activity patterns accompanied by nonlinear changes in network similarity profiles across levels of cognitive demand. Furthermore, the relative activation map-based whole-brain pattern analysis used in this study may provide a useful approach for evaluating changes in whole-brain state representations associated with cognitive load.

## Introduction

Working memory (WM) is a core cognitive function that supports attention, cognitive control, and goal-directed behavior. The N-back task is one of the most widely used experimental paradigms for investigating WM function, in which cognitive demand is assumed to increase parametrically with increasing memory load (Braver et al., 1997; Jonides et al., 1997; Owen et al., 2005). Consistent with this assumption, previous behavioral and neuroimaging studies have consistently reported load-dependent changes in both task performance and brain activity. Increased activation within the frontoparietal network (FPN), including the dorsolateral prefrontal cortex (DLPFC), and suppression of the default mode network (DMN) are well-established neural signatures of the N-back task (Owen et al., 2005; Arsalidou et al., 2013). These findings have generally been interpreted within a resource model framework, in which cognitive resources are progressively recruited as task demands increase. More recently, however, load-dependent changes have increasingly been conceptualized not simply as local increases in neural activity, but as dynamic reconfiguration of large-scale brain networks.

To date, WM has been investigated using a wide range of neuroimaging approaches, including regional activation analyses, functional connectivity analyses, multivariate decoding, and representational similarity analysis (RSA) (Braun et al., 2015; Shine et al., 2016; Harrison & Tong, 2009; Christophel et al., 2012; Skalaban et al., 2024). Collectively, these studies suggest that WM can be characterized not only in terms of activation magnitude, but also in terms of large-scale network organization and neural representational structure. Such findings highlight the importance of understanding WM processes beyond local activation changes and motivate whole-brain approaches that capture network-level and representational properties of brain activity. Recent studies have shown that large-scale brain networks, including the FPN, dorsal attention network (DAN), and DMN, exhibit dynamic changes in their organization during cognitive task performance (Vatansever et al., 2015; Shine et al., 2016; Zuo et al., 2018). Furthermore, flexible integration and segregation among networks are critical for cognitive control (Braun et al., 2015; Shine et al., 2016; Cohen & D’Esposito, 2016).

Although increasing WM load is generally associated with systematic changes in neural activity and large-scale network organization, these changes are not always monotonic. For example, prefrontal activity has been shown to increase from low to moderate loads but to exhibit inverted-U-like responses near capacity limits (Callicott et al., 1999; Cappell et al., 2010; Van Snellenberg et al., 2015). Moreover, individual differences and neural variability increase under high-load conditions, and load-dependent neural changes have been shown to strongly predict behavioral performance (Lamichhane et al., 2020).

Importantly, high-load conditions may induce distinct cognitive states, including capacity limitation, strategic adaptation, or disengagement from the task (Callicott et al., 1999; Todd & Marois, 2004; Van Snellenberg et al., 2015). Thus, high WM load may not simply reflect greater cognitive demand but may instead induce qualitative changes in cognitive processing. Accordingly, examining such high-load conditions may provide insight not only into stronger cognitive demands but also into qualitative changes in behavior and whole-brain activity patterns that emerge near working-memory capacity limits. Characterizing these changes may help clarify whether the increase in load primarily reflects a quantitative increase in processing demands or whether distinct cognitive processes become engaged under extreme task demands. However, it remains unclear whether increasing WM load primarily reflects a continuous modulation of cognitive processing or the emergence of qualitatively different whole-brain activity patterns under high-load conditions. Moreover, most previous N-back studies have focused on relatively limited load ranges (primarily 0–3-back), leaving neural representational changes under high-load conditions near or beyond WM capacity insufficiently characterized.

To better understand cognitive processing under high-load conditions, it may therefore be necessary to evaluate not only local activation changes but also the organization of whole-brain activity patterns themselves. Notably, all N-back conditions share common task components, including visual processing, motor responses, and sustained attention. Because these shared components contribute similarly across N-back conditions, condition-specific activation maps may be dominated by variance shared across task conditions. Consequently, load-specific differences may be more readily characterized by focusing on variation across load conditions. To address this issue, we focused on relative deviations from common task-related activity by computing relative activation maps, in which the participant-wise mean activation across all conditions was subtracted from each condition-specific activation map.

In the present study, we analyzed behavioral performance and fMRI activity across an extended 0–7-back paradigm to characterize load-dependent changes in whole-brain activity patterns across a broader range of WM demands than has typically been examined. A key question was whether increasing WM load produces continuous changes in whole-brain activity patterns or whether qualitatively different activity patterns emerge under high-load conditions. Specifically, we evaluated (i) load-dependent changes in whole-brain activity patterns, (ii) changes in similarity to large-scale brain networks, and (iii) changes in semantic representations. Through this approach, we aimed to characterize load-dependent changes in the N-back task from the perspective of whole-brain neural representations rather than regional activation magnitude alone.

## Methods

### 2.1 Experimental dataset

In the present study, we used a subset of the previously published HCP-mini/N-back EEG-fMRI dataset (Hiromitsu et al., 2026). In this dataset, EEG and fMRI sessions were conducted on separate days using a common task paradigm, and session order was counterbalanced across participants. The mean interval between sessions was 17.72 ± 10.94 days. The present study focused on fMRI data acquired during the extended N-back task (0–7-back) to examine load-dependent changes in whole-brain activity patterns associated with working memory demand. Participants consisted of 58 healthy adults (14 females; mean age = 23.07 ± 1.99 years). Written informed consent was obtained from all participants before participation. The study was approved by the ethics committee of the Advanced Telecommunications Research Institute International (ATR) and conducted in accordance with the Declaration of Helsinki.

The analyzed task consisted of a visual N-back paradigm comprising eight load conditions ranging from 0-back to 7-back (Fig. 1). The paradigm parametrically manipulated working memory load across a broader range than conventional N-back tasks. On each trial, a single alphabetic letter was presented, and participants judged whether the current stimulus matched the one presented N trials earlier. Each block consisted of 50 + N trials, while the proportions of target and lure trials were held constant across conditions. The task was designed such that the number of effective trials analyzed was equivalent across all conditions. Each load condition was presented once within each run, and the order of load conditions was randomized independently for each participant and run. Participants completed two runs, resulting in two repetitions of each load condition.

**Figure 1.**
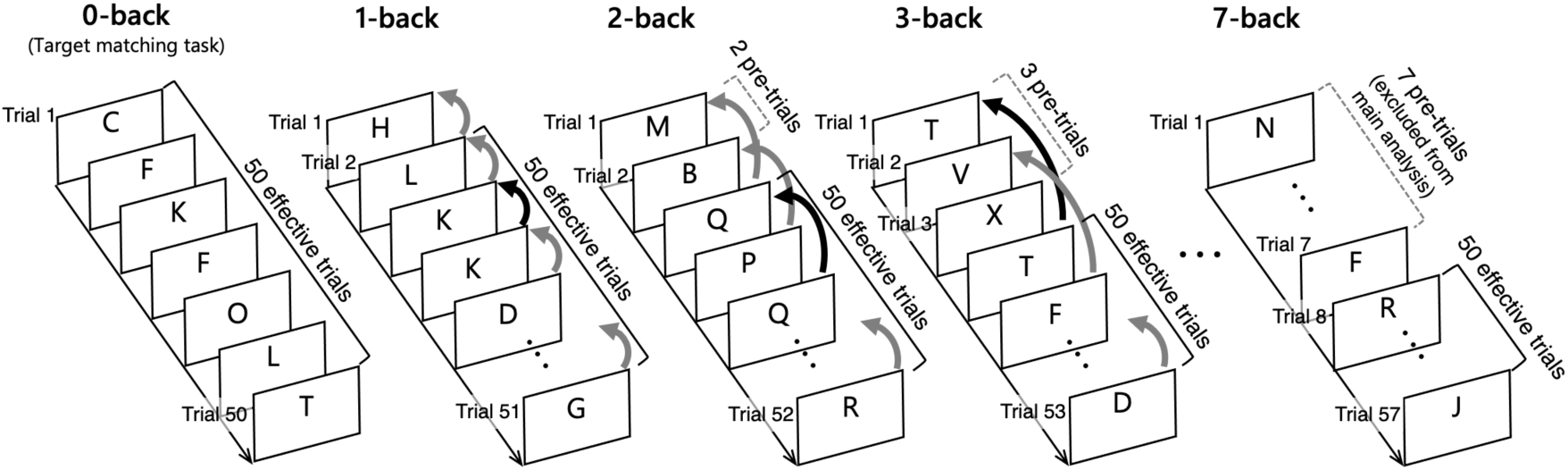
Schematic illustration of the extended N-back task. The task consisted of eight N-back-load conditions ranging from 0-back to 7-back. In the 0-back condition, participants performed a target-matching task, whereas in the N-back conditions (1–7-back), participants judged whether the current stimulus matched the stimulus presented N trials earlier. Each block consisted of 50 + N trials. The initial N pre-trials were excluded from the main analyses, resulting in 50 effective trials per condition.

MRI data were acquired using a 3T Siemens MAGNETOM Prisma scanner equipped with a 32-channel head coil. Functional images were obtained using a multiband echo-planar imaging (EPI) sequence (TR = 0.8 s, TE = 34 ms, voxel size = 2.4 mm isotropic, multiband factor = 6). Two functional runs were acquired with opposite phase-encoding directions: anterior-to-posterior (AP) and posterior-to-anterior (PA), with each load condition presented once in each run. Reverse phase-encoding images were also acquired for distortion correction. Structural images consisted of T1-weighted anatomical scans with 0.8-mm isotropic resolution. Detailed acquisition parameters and experimental procedures are described in Hiromitsu et al. (2026).

### 2.2 Behavioral measures

For each participant, discrimination sensitivity (d′), response bias (criterion C), and reaction time (RT) were calculated for each N-back condition based on signal detection theory. Because N-back judgments cannot be performed during the first N trials of each condition, these pre-trials were excluded from behavioral analyses. As a result, 50 effective trials per condition were analyzed (Fig. 1). d′ was used as a bias-free measure of discrimination performance and has been shown to provide a robust index of N-back task performance (Haatveit et al., 2010; Meule, 2017). Hit rates and false alarm rates were calculated for each condition, and d′ and criterion C were derived from these values. To avoid infinite values in d′ estimation, a log-linear correction (Hautus, 1995) was applied when hit rates or false alarm rates reached 0 or 1. Specifically, 0.5 was added to the number of hits and false alarms, and 1 was added to the number of signal and noise trials. Trials with anticipatory responses (<150 ms) or extremely long RT values (>2.5 standard deviations from the participant- and condition-specific mean) were excluded from RT analyses.

### 2.3 MRI preprocessing

fMRI preprocessing was performed using FSL and SPM12. Susceptibility distortion correction was first performed with TOPUP using reverse-phase-encoding images, followed by motion correction, spatial registration, and normalization to MNI standard space. Spatial smoothing was applied using a Gaussian kernel with a full width at half maximum (FWHM) of 6 mm. Following preprocessing, analyses were restricted to voxels within a gray matter mask. All analyses were conducted using normalized functional images in MNI space. Detailed preprocessing procedures are described in Hiromitsu et al. (2026).

### 2.4 fMRI analysis

First-level analyses were performed using a general linear model (GLM) implemented in SPM12. Each N-back condition (0–7-back) was modeled as a boxcar regressor corresponding to the duration of effective trials within each block and convolved with the canonical hemodynamic response function (HRF). The initial N pre-trials for each condition were modeled separately as independent regressors because N-back judgments were not yet available during these periods. Six motion parameters were included as nuisance regressors. To remove low-frequency drift while preserving task-related signals associated with the long block design (∼80 s), a high-pass filter cutoff of 256 s was used. Temporal autocorrelation was modeled using the first-order autoregressive model (AR (1)) implemented in SPM12.

### 2.5 Whole-brain pattern analysis

To characterize load-dependent whole-brain activity patterns, relative activation maps were computed for each condition (Fig. 2A). First, condition-specific activation maps corresponding to each N-back condition (0–7-back) were estimated from the first-level GLM for each participant. Inspired by previous task-battery studies emphasizing task-specific deviations from common activation patterns (King et al., 2019), the participant-wise mean activation map across all conditions was subtracted from each condition-specific activation map to obtain relative activation maps:

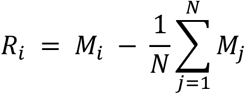

where (*Ri*) represents the relative activation map for condition (*i*), (*Mi*) represents the activation map for condition (*i*), and (*N*) denotes the total number of conditions. Subtracting the participant-wise mean activity pattern reduced activity components shared across conditions, such as visual processing, motor responses, and sustained attention demands, thereby emphasizing load-specific spatial activity configurations. Consequently, the analysis focused on relative differences in spatial activity patterns rather than absolute signal magnitude. This approach also enabled direct comparison across all load conditions without requiring a specific baseline condition.

For group-level visualization, relative activation maps for each condition were entered into second-level random-effects analyses to generate group-level statistical maps (t-maps) (Fig. 2A). To compare relative spatial distributions across conditions, voxel values within the gray matter mask were z-score normalized.

**Figure 2.**
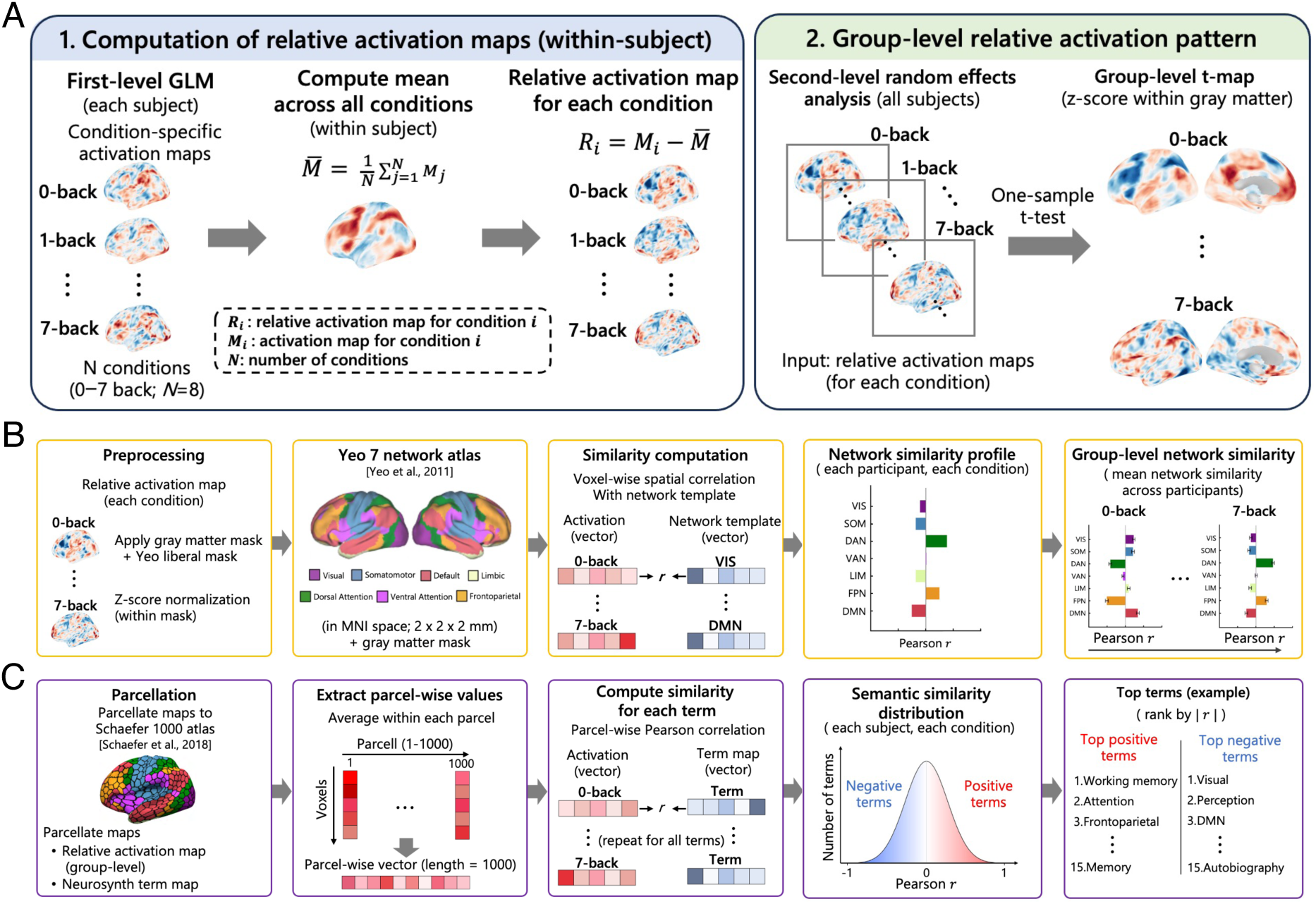
Overview of the analysis pipeline for relative activation map–based whole-brain pattern analysis. (A) Computation of relative activation maps within each participant. Condition-specific activation maps obtained from the first-level GLM were averaged across all N-back conditions (0–7-back), and the mean activation map was subtracted from each condition map to obtain relative activation maps that represent condition-specific deviations from the participant’s average task-related activity pattern. Group-level relative activation maps were then estimated using second-level random-effects analysis. (B) Network similarity analysis using the Yeo 7-network atlas. Relative activation maps were masked with the gray matter and Yeo liberal masks, z-score-normalized within each mask, and spatially correlated with each network template across gray matter voxels. This analysis yielded condition-specific network similarity profiles for each participant and group-level mean similarity patterns across N-back loads. (C) Semantic similarity analysis using Neurosynth-derived coordinate-count term maps. Relative activation maps and Neurosynth-derived semantic maps were parcellated using the Schaefer 1000-parcel atlas, and parcel-wise similarity was computed using Pearson correlation. The resulting semantic similarity distributions were used to identify cognitive terms showing positive or negative associations with each load condition.

### 2.6 Pattern similarity analysis

To evaluate the extent to which whole-brain activity patterns resembled canonical large-scale brain networks, network similarity analyses were performed on first-level relative activation maps. Spatial similarity between condition-specific activity patterns and the Yeo 7-network atlas (Yeo et al., 2011) was calculated (Fig. 2B). All activation maps and network templates were resliced into a common MNI space (2 × 2 × 2 mm). Network templates were masked with a gray matter mask. The Yeo network templates were binary masks, with voxels belonging to a given network assigned a value of 1 and all other voxels assigned a value of 0. Activation maps were masked with both the gray matter mask and the Yeo liberal mask and subsequently z-scored within the masked voxels.

For each participant and condition, voxel-wise Pearson correlation coefficients were calculated between activity patterns and each network template, yielding condition-specific network similarity profiles. Correlation coefficients were subsequently averaged across participants to obtain group-level similarity measures.

### 2.7 Semantic similarity analyses

Semantic similarity analyses were additionally performed using Neurosynth-derived semantic maps based on coordinate-count distributions (Yarkoni et al., 2011) to characterize cognitive representations associated with each load condition (Fig. 2C). We constructed coordinate-count maps for individual Neurosynth terms using the coordinate and annotation files obtained through the NiMARE library. The coordinate file contained reported stereotactic activation foci for each study, and the annotation file contained term frequency-inverse document frequency (TF–IDF) scores derived from article abstracts for a large vocabulary of terms. Study identifiers were converted to character strings before matching the two tables. For each term, studies were selected if their TF–IDF score for that term was greater than zero. Terms associated with fewer than 10 studies were excluded. For each remaining term, all reported foci from the selected studies were extracted. The coordinates were then discretized onto a 2-mm MNI152 grid with dimensions 91 × 109 × 91. Only foci falling within the grid were retained. For each term, a three-dimensional count image was created by adding one count to the voxel corresponding to each retained focus. The resulting maps, therefore, reflected the spatial distribution of reported activation foci from studies in which the term appeared in the abstract-derived Neurosynth annotation (Asai et al., 2026).

Importantly, these images should be interpreted as term-associated coordinate-count maps rather than as formal Neurosynth statistical association maps. The term-wise count maps were saved as NIfTI images and subsequently summarized within the Schaefer 1000-parcel cortical atlas (Schaefer et al., 2018). For each map, voxel values were averaged within each Schaefer parcel, yielding a 1000 × N term matrix whose entries represented the mean coordinate-count density for each term within each parcel. This matrix was used as the parcellated representation of Neurosynth-derived semantic brain maps.

Semantic similarity analyses were performed using group-level relative activation maps obtained from the second-level random-effects analysis. Relative activation maps were parcellated using the same atlas to ensure spatial correspondence between datasets. Parcel-wise Pearson correlations were then computed between each condition-specific activity pattern and each Neurosynth-derived semantic map. Terms were ranked according to correlation magnitude, and the top positively and negatively associated terms were extracted for visualization and interpretation. Because relative activation maps reflect deviations from participant-wise mean activity patterns, this analysis evaluated relative differences in cognitive representations across load conditions.

### 2.8 Statistical analysis

Behavioral measures (d′, criterion C, and RT) and network similarity measures were analyzed using repeated-measures analyses of variance (ANOVAs) with N-back load condition (0–7-back) as a within-subject factor. Greenhouse–Geisser correction was applied when necessary. To assess both linear and nonlinear (inverted-U–like) load-dependent changes, linear mixed-effects (LME) models including linear and quadratic load terms were additionally applied. Models included N-back load (linear and quadratic terms) as fixed effects and participant as a random effect. For criterion C, an additional model including d′ as a covariate was also examined. Bonferroni correction was applied for post hoc comparisons, and false discovery rate (FDR) correction was applied for multiple-network analyses.

## Results

### 3.1 Behavioral performance

Behavioral performance exhibited systematic load-dependent changes across N-back conditions (Fig. 3). Discrimination sensitivity (d′) progressively declined with increasing WM load, indicating a marked performance decline under high-load conditions. In contrast, criterion C gradually increased with load, reflecting increasingly conservative response tendencies under higher cognitive demand. RT exhibited an inverted-U profile, peaking at intermediate-load conditions.

**Figure 3.**
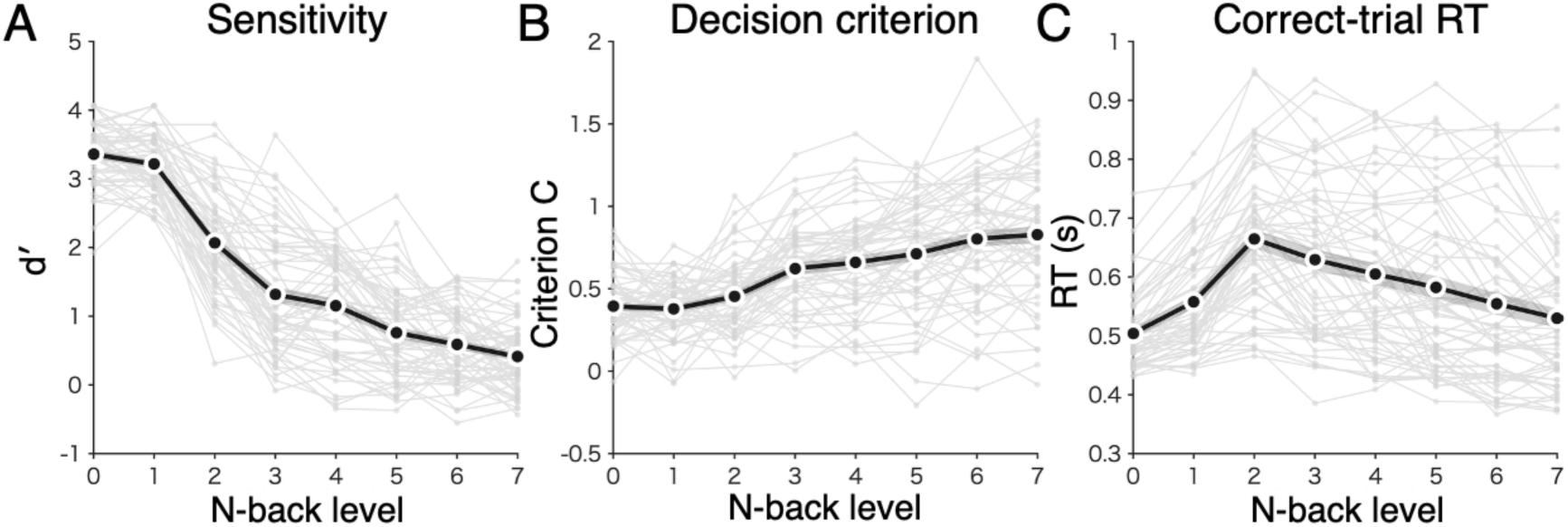
Behavioral performance across working memory load. (A) Sensitivity (d′), (B) Response bias (criterion C), and (C) correct-trial reaction time (RT) across N-back load conditions (0–7-back). Gray lines indicate individual participants, and black lines indicate group means. Sensitivity progressively decreased across N-back load conditions, whereas criterion C gradually increased, indicating a more conservative response tendency at higher loads. Reaction time exhibited an inverted-U pattern, peaking at intermediate load conditions.

Repeated-measures ANOVA revealed significant main effects of N-back load on d′ [F (4.49, 256.09) = 365.17, *p* < .001], criterion C [F (3.58, 204.30) = 40.00, *p* < .001], and RT [F (3.34, 190.62) = 36.55, *p* < .001]. To further characterize the shapes of these load-dependent changes, LME models including both linear and quadratic terms were applied.

For d′, both significant linear and quadratic effects were observed (linear: β = −0.84, p < .001; quadratic: β = 0.06, *p* < .001), indicating a nonlinear decline in discrimination sensitivity with increasing load. For criterion C, a significant positive linear effect was observed (linear: β = 0.08, *p* < .001), whereas the quadratic effect was not significant (quadratic: β = −0.001, *p* > .05). However, when d′ was included as a covariate, the linear effect was no longer significant and only the quadratic effect remained (linear: β = −0.008, *p* > .05; quadratic: β = 0.005, *p* < .05; d′: β = −0.103, *p* < .001). These findings suggest that load-dependent changes in criterion C were largely attributable to reductions in discrimination sensitivity, while also containing variance not fully explained by d′ alone.

Similarly, RT showed significant linear and quadratic effects (linear: β = 0.063, p < .001; quadratic: β = −0.009, p < .001), confirming an inverted-U pattern with maximal RT observed at intermediate load conditions.

### 3.2 Load-dependent changes in whole-brain activity patterns

To evaluate load-dependent changes in whole-brain activity patterns, group-level relative activation maps were visualized for each N-back condition (Fig. 4A). Low-load conditions (0–1-back) were characterized by relative activity patterns centered on DMN-related regions. In contrast, intermediate-load conditions (2–4-back) showed relative activity patterns dominated by DAN- and FPN-related regions. Under high-load conditions, the spatial distribution of activity patterns shifted, exhibiting configurations distinct from those observed under low- and intermediate-load conditions.

**Figure 4.**
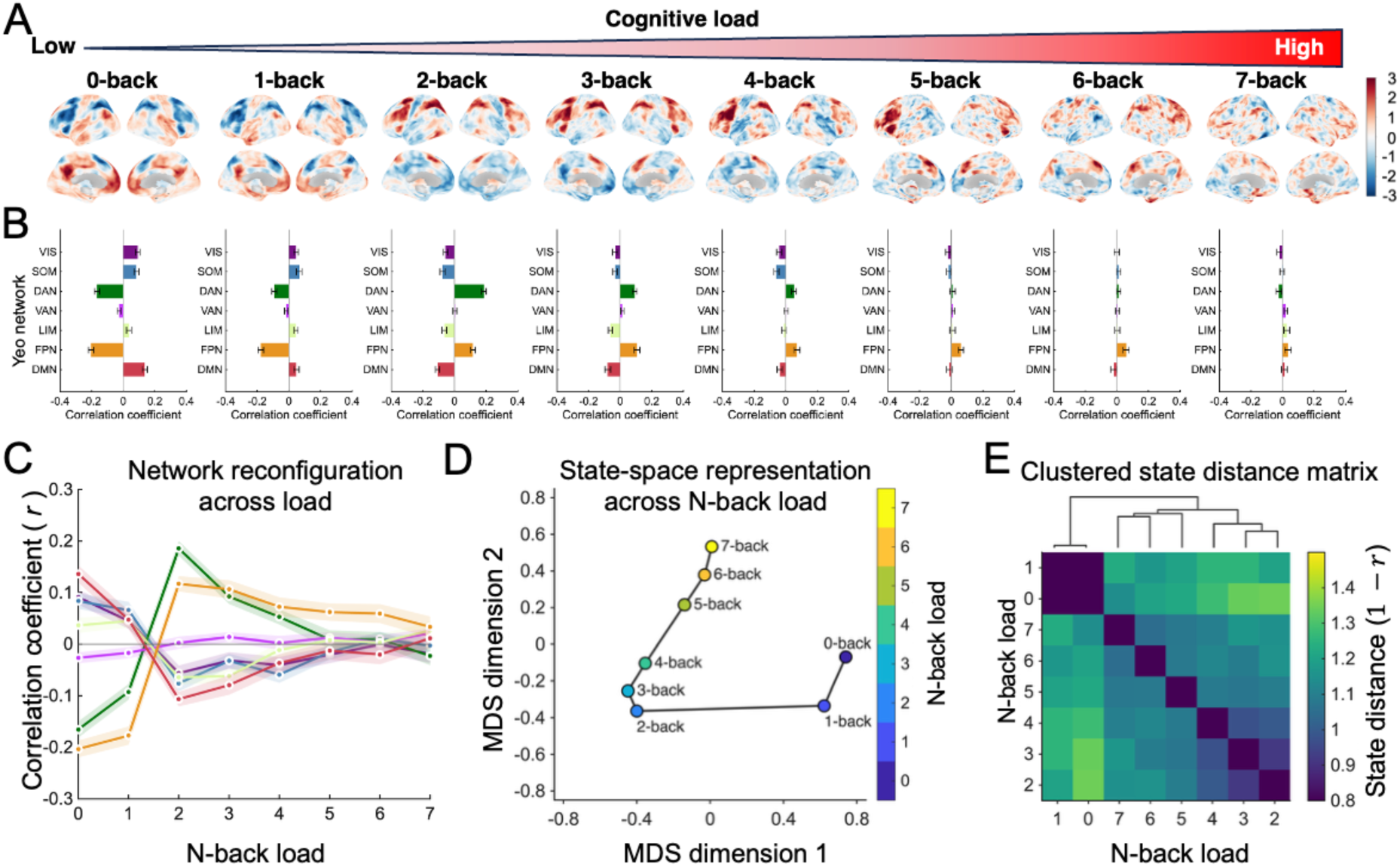
Load-dependent changes in whole-brain activity patterns and network similarity. (A) Group-level relative activation maps across N-back load conditions (0–7-back). Low-load conditions showed relative activity patterns centered on default mode regions, whereas intermediate-load conditions showed frontoparietal-dominant patterns. High-load conditions exhibited altered whole-brain spatial configurations. (B) Network-level similarity profiles between relative activation maps and Yeo 7-network templates for each load condition. (C) Load-dependent changes in network similarity. DAN and FPN similarity exhibited inverted-U–like patterns peaking at intermediate loads, whereas DMN similarity decreased at intermediate loads and partially recovered at high loads. Error bars indicate SEM across participants. (D) State-space representation of load-dependent activity patterns. (E) Distance matrix and hierarchical clustering of group-level relative activation maps across load conditions, showing separation between low-load and higher-load states.

To further visualize relationships among load conditions, pairwise distances between condition-specific relative activation maps were computed for each participant and then averaged across participants. Multidimensional scaling (MDS) was subsequently applied to the resulting group-average distance matrix (Fig. 4D). The 0-back and 1-back conditions formed a closely clustered group, while conditions from 2-back onward occupied increasingly separated positions in the low-dimensional state space. Consistent with the MDS results, the group-average distance matrix and hierarchical clustering analysis demonstrated a separation between low-load conditions (0–1-back) and higher-load conditions (2–7-back) (Fig. 4E).

### 3.3 Network-level changes across working memory load

To quantify the network-level organization of whole-brain activity patterns, spatial similarity between relative activation maps and the Yeo 7-network templates was calculated (Fig. 4B). Distinct network similarity profiles were observed across low-, intermediate-, and high-load conditions.

In particular, the dorsal attention network (DAN) similarity exhibited an inverted-U profile, peaking around the 2-back condition. In contrast, similarity to the default mode network (DMN) was highest under low-load conditions, reached a minimum around 2-back, and partially recovered under high-load conditions. Similarity to the frontoparietal network (FPN) also peaked highest around 2-back and gradually decreased thereafter, although FPN similarity remained relatively preserved compared with other networks.

Repeated-measures ANOVA revealed significant load effects for VIS [*F*(7, 399) = 8.228, FDR-corrected *p* < .001], SOM [*F*(7, 399) = 12.524, FDR-corrected *p* < .001], DAN [*F*(7, 399) = 43.062, FDR-corrected *p* < .001], LIM [*F*(7, 399) = 6.670, FDR-corrected *p* < .001], FPN [*F*(5.56, 316.96) = 44.736, FDR-corrected *p* < .001], and DMN [*F*(7, 399) = 16.366, FDR-corrected *p* < .001], whereas no significant load effect was observed for VAN [*F*(5.69, 324.14) = 1.708, FDR-corrected *p* > .05].

LME analyses further demonstrated significant positive linear and negative quadratic effects for DAN similarity (linear: *β* = 0.027, FDR-corrected *p* < .001; quadratic: *β* = −0.097, FDR-corrected *p* < .001), confirming an inverted-U profile peaking at intermediate load conditions. Similar effects were observed for FPN similarity (linear: *β* = 0.077, FDR-corrected *p* < .001; quadratic: *β* = −0.098, FDR-corrected *p* < .001). In contrast, DMN similarity showed a significant negative linear effect and positive quadratic effect (linear: β = −0.026, FDR-corrected *p* < .001; quadratic: β = 0.071, FDR-corrected *p* < .001), reflecting initial suppression followed by partial recovery under high-load conditions (Fig. 4C).

### 3.4 Load-dependent changes in semantic representations

To investigate load-dependent changes in cognitive representations, spatial similarity between condition-specific relative activation maps and Neurosynth-derived semantic maps was calculated (Fig. 5). Distinct semantic representation profiles emerged across load conditions.

**Figure 5.**
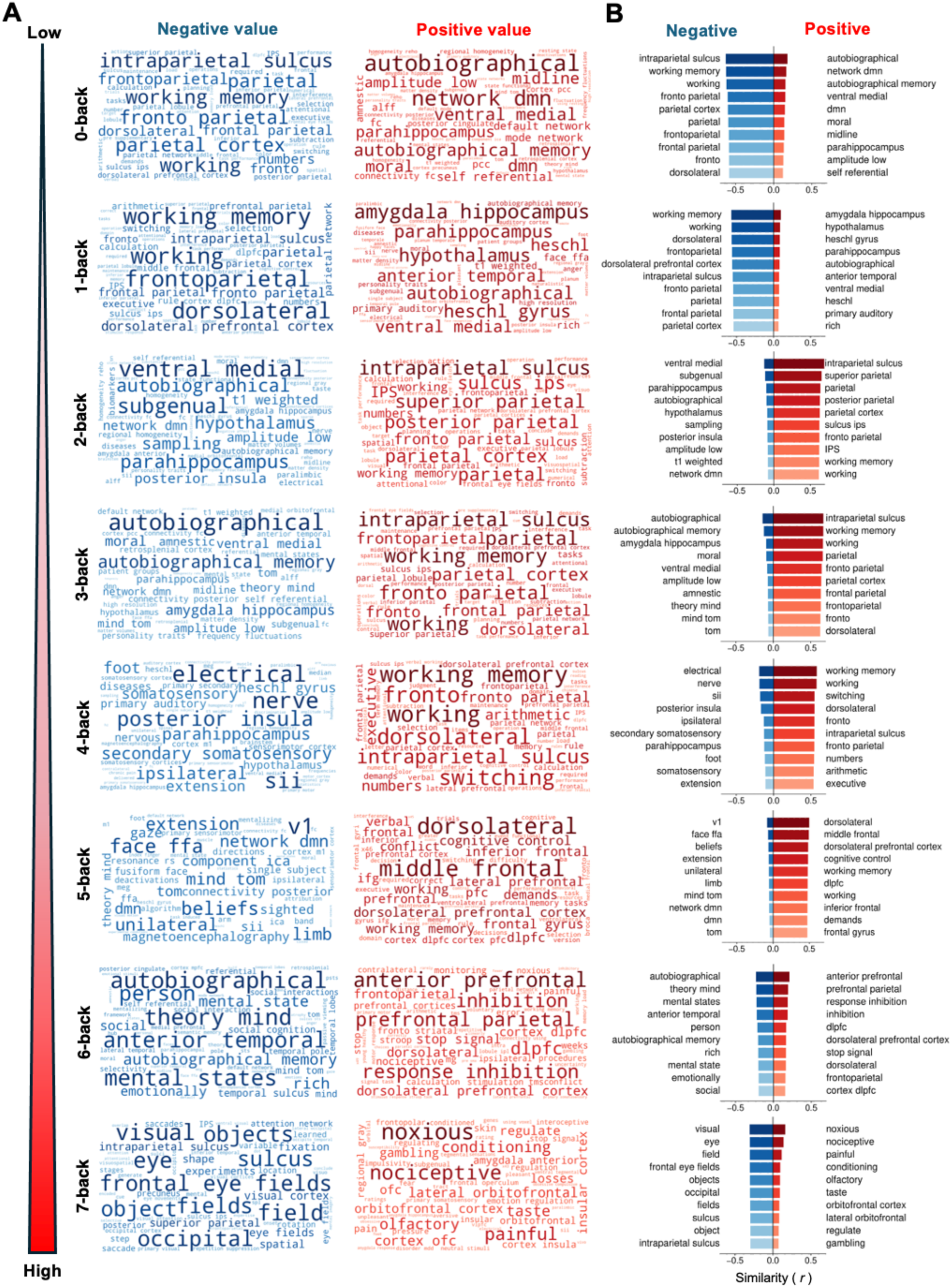
Load-dependent changes in semantic representations across N-back conditions. (A) Word clouds showing the most negatively (blue) and positively (red) associated Neurosynth terms for the relative activation map of each N-back condition. Term size reflects the relative strength of association between the condition-specific relative activation map and Neurosynth-derived semantic maps. (B) Top 10 positively and negatively associated Neurosynth terms ranked by spatial similarity for each load condition. Low-load conditions (0–1-back) were characterized by default mode, autobiographical memory, and self-referential representations. Intermediate-load conditions (2–5-back) showed prominent associations with working memory, frontoparietal, and attention-related representations. High-load conditions (6–7-back) exhibited increased similarity to inhibition-, salience-, and aversive/interoceptive-related representations.

Low-load conditions showed relatively high similarity to terms associated with autobiographical memory, self-referential processing, and default mode network function. Intermediate-load conditions (2–5-back) exhibited increased similarity to terms related to working memory, cognitive control, DLPFC function, and frontoparietal processing. In contrast, the 6-back condition showed stronger associations with response inhibition-related terms. The 7-back condition, meanwhile, exhibited increased similarity to terms related to salience and aversive/interoceptive processing, including noxious, nociceptive, insular cortex, and anterior amygdala.

## Discussion

Collectively, the present findings demonstrate load-dependent changes in whole-brain activity patterns across an extended range of working-memory demands. Behavioral performance and network similarity profiles exhibited nonlinear load-dependent changes, suggesting that increasing WM load cannot be fully characterized as a simple monotonic increase in cognitive demand. Using an extended N-back paradigm spanning 0-back to 7-back conditions, we observed convergent changes across behavioral performance, network similarity profiles, and semantic representations. DAN and FPN similarity exhibited inverted-U profiles peaking under intermediate-load conditions. At high-load conditions, DMN similarity partially recovered together with increased salience-, aversive/interoceptive-, and inhibitory-control-related representations. These findings suggest that increasing WM load does not simply produce continuous escalation of cognitive demand but is associated with shifts in whole-brain activity patterns across load conditions. In particular, the results suggest that high-load conditions may involve cognitive processes that differ from those engaged at intermediate load levels.

### 4.1 Intermediate-load task-engaged configuration

Behavioral performance and large-scale network organization both exhibited characteristic changes under intermediate-load conditions, particularly around the 2-back condition. RT reached its maximum at intermediate loads, accompanied by peak DAN and FPN similarity. Semantic similarity analyses further revealed increased similarity to working memory-, frontoparietal control-, and executive control-related representations during these conditions. Together, these findings suggest that intermediate-load conditions may correspond to a maximally task-engaged configuration involving attentional allocation, cognitive control, and WM maintenance processes.

This interpretation is broadly consistent with previous studies reporting that frontoparietal activity does not increase monotonically with WM load but instead exhibits inverted-U-shaped responses across increasing levels of WM load (Callicott et al., 1999; Van Snellenberg et al., 2015). Under high-load conditions, frontoparietal activity typically falls along the descending portion of this profile. Lamichhane et al. (2020) similarly demonstrated that frontoparietal activity can decrease under high-load conditions and that such changes are associated with behavioral performance. Accordingly, the DAN and FPN similarity peaks observed in the present study may reflect not merely increased activation magnitude, but rather activity patterns associated with task-focused attentional and control processes that support optimal task performance.

Notably, the DAN is strongly implicated in goal-directed attention and externally oriented attentional control (Corbetta & Shulman, 2002). The present finding that DAN similarity peaked under intermediate-load conditions suggests that attentional control demands may be insufficient at low-load conditions. At high-load conditions, task demands may exceed processing capacity, making efficient externally directed control states difficult to sustain.

In addition, portions of the frontoparietal cortex have been conceptualized as belonging to the multiple-demand (MD) system, which is consistently recruited across diverse cognitive tasks, including working memory, task switching, and inhibitory control (Cole & Schneider, 2007; Duncan, 2010; Gratton et al., 2018; Assem et al., 2020; Duncan et al., 2020). Beyond reflecting cognitive effort, the MD system has also been shown to encode multiple forms of task-relevant information, including stimuli, rules, and responses (Duncan, 2001; Woolgar et al., 2016; Assem et al., 2020; Zheng et al., 2024). Increased FPN similarity under intermediate-load conditions may therefore reflect greater involvement of task-rule maintenance and goal-directed control processes within a stable task-engaged configuration.

### 4.2 Beyond capacity: overload-related changes in activity patterns

High-load conditions exhibited whole-brain activity patterns distinct from those observed under intermediate-load conditions. DAN and FPN similarity gradually declined beyond intermediate-load conditions, whereas DMN similarity partially recovered under high-load conditions. Semantic similarity analyses additionally revealed reduced dominance of working memory- and frontoparietal control-related terms together with increased similarity to salience-, aversive/interoceptive-, and inhibitory-control-related representations.

These findings suggest that high-load conditions may mark a shift from increasing WM demand to broader changes in whole-brain activity patterns. Previous studies have similarly reported plateauing or reduced frontoparietal activity near WM capacity limits (Lamichhane et al., 2020). The observed reduction in DAN and FPN similarity may therefore reflect difficulty maintaining task-focused attentional and control states once task demands exceed processing capacity, resulting in increasingly unstable and heterogeneous activity patterns.

Importantly, the partial recovery of DMN similarity under high-load conditions does not necessarily indicate simple task disengagement. The DMN has increasingly been reconceptualized not merely as a task-negative network, but as a core network involved in memory, self-related processing, and conceptual cognition (Menon, 2023). Furthermore, DMN connectivity dynamically changes according to WM demands (Vatansever et al., 2015). Accordingly, the observed recovery of DMN similarity may reflect increased internally oriented processing or a rebalancing of activity between FPN- and DMN-related systems.

Taken together, these findings suggest that high-load conditions are associated with activity patterns that differ from those observed under intermediate-load conditions and may reflect processing associated with capacity limitations, involving salience/interoceptive systems and inhibitory-control-related processes. This interpretation is conceptually related to previous accounts of working-memory overload (Yun et al., 2010). However, the present findings alone do not allow direct inference about specific psychological states. Rather, the results are more appropriately interpreted as reflecting a shift from task-engaged activity patterns toward alternative whole-brain activity patterns characterized by greater involvement of salience-related systems and DMN-associated processing.

### 4.3 Network-level changes under N-back load

The present findings are broadly consistent with contemporary network neuroscience perspectives emphasizing the involvement of large-scale brain networks in working memory performance. Previous studies have demonstrated flexible interactions among the FPN, DAN, DMN, and salience network depending on cognitive demand and task requirements (Braun et al., 2015; Shine et al., 2016; Liang et al., 2016; Causse et al., 2022). Consistent with these findings, DAN and FPN similarity was highest under intermediate-load conditions, indicating that activity patterns during these conditions more closely resembled DAN- and FPN-related network templates.

Liang et al. (2016) further demonstrated that the salience network dynamically alters its connectivity with both the DMN and executive control network as WM load increases. The increased salience-related semantic representations observed under high-load conditions are consistent with the possibility that salience-related systems become increasingly engaged during overload states, potentially influencing interactions between the FPN and DMN.

### 4.4 Methodological implications of whole-brain pattern analysis

Unlike conventional region-based activation analyses, the present study employed whole-brain pattern analysis using relative activation maps. Previous N-back studies have typically investigated load-dependent brain activity using regional activation analysis, functional connectivity analysis, representational similarity analysis (RSA), or decoding approaches. In contrast, the present framework treated each load condition as a whole-brain activity configuration and evaluated its network organization and semantic representation in an integrated manner.

Because stimulus structure and task format were relatively consistent across N-back conditions, relative activation maps facilitated characterization of load-specific large-scale activity configurations by reducing the contribution of processes shared across conditions, including visual input, motor responses, and sustained attention. This framework enabled comparison across the broad 0–7-back load range within a common representational space without relying solely on absolute activation increases or pairwise condition contrasts.

Importantly, the DAN/FPN/DMN patterns observed in the present study were broadly consistent with findings from regional activation analyses, functional connectivity studies, and studies of brain dynamics during working memory tasks. Specifically, intermediate-load conditions showed maximal DAN/FPN similarity, whereas high-load conditions were associated with reduced DAN/FPN similarity and partial recovery of DMN similarity. These patterns are broadly compatible with previous reports of maximal task-positive network engagement at moderate loads and reduced frontoparietal recruitment under high-load conditions. These findings suggest that relative whole-brain pattern analysis preserves key properties of previously reported network-level findings while additionally enabling integrated characterization of large-scale representational organization.

It is also important to note that relative activation maps evaluate relative activity configurations rather than absolute activation magnitude. Consequently, reductions in DAN and FPN similarity under high-load conditions should not necessarily be interpreted as a simple reduction in activation. Rather, they may indicate a change in task-engaged activity patterns together with increasing interindividual heterogeneity in large-scale brain state.

### 4.5 Implications for extended N-back paradigms

A major strength of the present study is the use of an extended load range spanning 0–7-back conditions, which enabled characterization of neural activity across a substantially broader range of cognitive demand than has typically been examined. The extended load range revealed changes in whole-brain activity patterns across load conditions, suggesting that increasing working-memory demands may be accompanied by qualitative changes in cognitive state rather than merely quantitative increases in processing load.

Although the N-back task is a central paradigm for investigating working memory and executive function, previous studies have shown that observed neural responses and their interpretations can vary substantially depending on task structure, load range, and analytical approach (Huang et al., 2025). In the present paradigm, N-back load varied dynamically within each run, requiring participants to flexibly adjust updating, maintenance, comparison, and response inhibition strategies according to the current load condition. Such load variability may impose greater demands on flexible cognitive control and strategic adjustment compared with paradigms using fixed-load block designs. This flexibility may reflect strategic adaptation to increasing working memory load and task difficulty during N-back performance. Accordingly, the inverted-U DAN/FPN profiles and increased salience-related representations observed under high-load conditions may reflect not only quantitative load effects, but also strategic adaptation to dynamically changing task demands.

### 4.6 Limitations

Several limitations should be acknowledged. First, because relative activation maps represent deviations from participant-wise mean activity patterns, reductions in network similarity under high-load conditions cannot fully distinguish between shared state changes across participants and cancellation effects arising from interindividual variability in peak load levels or cognitive strategies. Previous studies have suggested that inverted-U frontoparietal responses and peak positions may vary according to individual WM capacity and behavioral performance (Lamichhane et al., 2020). Future studies should therefore directly examine individual state trajectories and peak-load positions to better characterize cognitive-state diversification under high-load conditions.

Second, the semantic similarity analysis employed in the present study provides an indirect meta-analytic measure and should therefore be interpreted cautiously. Increased similarity to pain- or salience-related representations should not be taken as direct evidence of corresponding psychological states. Rather, these findings indicate that whole-brain activity patterns under high-load conditions exhibited spatial similarity to networks associated with cognitive, affective, and control-related processes represented in the Neurosynth database. Future studies incorporating subjective measures of effort, fatigue, mind-wandering, or perceived task difficulty, together with physiological measures, may help clarify the psychological significance of these semantic representations.

Third, the present analyses were based on condition-level activity patterns rather than network interactions or state-transition dynamics. Therefore, the observed changes should not be interpreted as direct evidence of large-scale network reconfiguration. In addition, the present analyses could not directly assess how brain states transitioned over time as task demands varied across the 0–7-back load range. Recent studies have increasingly emphasized the importance of dynamic analyses of large-scale network organization and state transitions, demonstrating relationships between dynamic integration/segregation processes and behavior during learning and cognitive tasks (Shine et al., 2016; Zuo et al., 2018; Wang et al., 2024). Future studies combining EEG and brain-dynamics analyses may therefore provide deeper insights into temporal changes in whole-brain configurations under high-load conditions.

## Conclusion

Using an extended N-back paradigm spanning 0–7-back conditions, the present study characterized load-dependent changes in whole-brain activity patterns associated with working memory demand. Across behavioral performance, network similarity profiles, and semantic representations, we observed convergent changes as load increased. Intermediate-load conditions were characterized by activity patterns associated with task-focused attentional and control processes. In contrast, high-load conditions showed distinct activity patterns together with increased associations with salience-, aversive/interoceptive-, and inhibitory-control-related representations. These findings demonstrate that extending the N-back paradigm beyond conventional load ranges can reveal changes in large-scale activity patterns that are not apparent at lower load. The relative whole-brain pattern analysis framework may provide a useful approach for investigating load-dependent changes in whole-brain activity patterns across a broad range of cognitive demands.

## Acknowledgments

During the preparation of this manuscript, the authors used ChatGPT (OpenAI; versions 5.4 and 5.5) to assist with literature review, programming, conceptual organization, and English-language editing. All scientific interpretations and final manuscript decisions were made by the authors.

